# Necessity and contingency in developmental genetic screens: LIN-3, Wnt and semaphorin pathways in vulval induction of the nematode *Oscheius tipulae*

**DOI:** 10.1101/383729

**Authors:** Amhed M. Vargas-Velazquez, Fabrice Besnard, Marie-Anne Félix

## Abstract

**Abstract:** Genetic screens in the nematode *Caenorhabditis elegans* identified the EGF/Ras and Notch pathways as central for vulval precursor cell fate patterning. Schematically, the anchor cell secretes EGF, inducing the P6.p cell to a 1° vulval fate; P6.p in turn induces its neighbors to a 2° fate through Delta-Notch signaling and represses Ras signaling. In the nematode *Oscheius tipulae*, the anchor cell successively induces 2° then 1° vulval fates. Here we report on the molecular identification of mutations affecting vulval induction in *O. tipulae*. A single Induction Vulvaless mutation was found, which we identify as a cis-regulatory deletion in a tissue-specific enhancer of the *O. tipulae lin-3* homolog, confirmed by CRISPR/Cas9 mutation. In contrast to this predictable Vulvaless mutation, mutations resulting in an excess of 2° fates unexpectedly correspond to the plexin/semaphorin pathway, which was not implicated in vulval fate induction in *C. elegans*. Hyperinduction of P4.p and P8.p in these mutants likely results from mispositioning of these cells due to a lack of contact inhibition. The third signaling pathway found by forward genetics in *O. tipulae* is the Wnt pathway: decrease in Wnt pathway activity results in loss of vulval precursor competence and induction, and 1° fate miscentering on P5.p. Our results suggest that the EGF and Wnt pathways have qualitatively similar activities in vulval induction in *C. elegans* and *O. tipulae*, albeit with quantitative differences in the effects of mutation. This study highlights both necessity and contingency in forward genetic screens.

**100-word summary:** Genetic screens in the nematode *Caenorhabditis elegans* identified EGF and Notch pathways as key for vulval precursor cell fate patterning. Here we report on the molecular identification of mutations affecting vulval induction in another nematode, *Oscheius tipulae*. The single mutation with reduced induction is identified as a *cis*-regulatory deletion in the *O. tipulae lin-3* homolog, confirmed by CRISPR/Cas9 mutation. In contrast to this predictable Vulvaless mutation, mutations resulting in an excess of 2° vulval fates unexpectedly correspond to the plexin/semaphorin pathway, not implicated in vulval induction in *C. elegans*. This study highlights both necessity and contingency in forward genetic screens.

## Introduction

How multicellular organisms arise from single cells is a question that has intrigued scientists over ages. In the 1960s, Sydney Brenner selected *Caenorhabditis elegans* as a new model organism to study animal development using genetics (Brenner 1974). Vulva precursor cell fate patterning rapidly became one of the most studied developmental processes in *C. elegans*, due to the easy isolation of mutants with a defective vulva (Sternberg 2005).

The *C. elegans* vulva is an epidermal specialization that develops from a row of six vulva precursor cells (VPCs) in the ventral epidermis, called P3.p to P8.p from anterior to posterior. In most animals, the central vulval fate, or 1° fate, is adopted by P6.p, while the outer vulval fate, or 2° fate, is adopted by its neighbors P5.p and P7.p (Sulston and Horvitz 1977; Sternberg 2005). Finally, P3.p, P4.p and P8.p are able to replace the central cells (for example if they are destroyed with a laser), but normally adopt a standard epidermal fate with one division and fusion of the daughters to the large epidermal syncytium hyp7 (Sulston and White 1980). Laser ablation of the anchor cell (AC) in the gonad primordium results in all precursor cells adopting a 3° fate, showing that the vulval fates are induced by the anchor cell (Kimble 1981).

Upon random chemical mutagenesis, some recurrent phenotypes were isolated with pronounced defects in vulva development, such as the Vul (Vulvaless) and Muv (Multivulva) phenotypes (Horvitz and Sulston 1980; Ferguson and Horvitz 1985). The Vulvaless mutants can be easily seen in the dissecting microscope by the internal hatching of the progeny in their mother (bag of worms). The Vulvaless mutants can be further classified in two classes, i) those that mimicked an AC ablation (cells adopting a 3° fate), or Induction Vulvaless and ii) those that prevented the development of competent vulva precursor cells, or Generation Vulvaless (Ferguson et al. 1987). The Multivulva mutants are recognized by the additional protrusions on the ventral cuticle (pseudovulvae).

The *C. elegans* Induction Vulvaless and the Multivulva mutants allowed the identification of the EGF/Ras/MAP kinase pathway, the former class corresponding to a loss of activity in the pathway, the latter to a gain of activity (Sternberg 2005). In addition, mutations at the *lin-12* locus affected 2° fates specifically: reduction-of-function *lin-12* alleles transformed 2° fates to 1° or 3°, while gain-of-function alleles transformed 1° and 3° fates to the 2° fate (Greenwald et al. 1983). *lin-12* was shown to encode a Notch receptor, receiving Delta signals mostly produced by P6.p. Studies of the interplay between the EGF and Delta/Notch pathways in patterning vulval cell fates established this system as a textbook example of intercellular signalling and organogenesis (Sternberg and Han 1998).

Since the 1990s, studies of vulva development in different *Caenorhabditis* species and other nematode genera have made vulva development an emblematic example of developmental system drift (DSD; True and Haag 2001): while the vulval cell fate pattern remains overall invariant (2°1°2° for P5.p, P6.p and P7.p), evolution occurs in the manner in which it forms. First, the size of the competence group varies (Sternberg and Horvitz 1982; Sommer and Sternberg 1996; Félix et al. 2000a; Delattre and Félix 2001; Pénigault and Félix 2011a). Second, vulval cell fate patterning does not always require the anchor cell (Sommer and Sternberg 1994; Félix et al. 2000a). Third, when it requires the gonad, ablating the anchor cell at intermediate timepoints has widely different effects depending on the species (Sommer and Sternberg 1994; Félix and Sternberg 1997; Sommer 2005; Kiontke et al. 2007; Félix and Barkoulas 2012; Félix 2012). Especially, in many genera of rhabditids and diplogastrids (Félix and Sternberg 1997; Félix and Sternberg 1998; Sigrist and Sommer 1999; Félix et al. 2000a; Félix 2007; Kiontke et al. 2007), the ablation at an intermediate timepoint results in P(5-7).p adopting a 2° fate (vs. a 3° fate for the outer cells), with no apparent differences among these three cells. This contrast with anchor cell ablation results in *C. elegans*, where no such intermediate state exists and P6.p adopts a 1° fate earlier, thereby activating lateral induction and inhibition (Félix 2007) (Fig. 1). The mode of induction where an intermediate fate is found for all cells has been called a two-step induction (Félix and Sternberg 1997). In this case, the second step of induction of the 1° fate occurs after one division round, on P6.p daughters. Signaling however may be continuous (Félix and Sternberg 1997; Sigrist and Sommer 1999; Kiontke et al. 2007).

**Figure 1.**
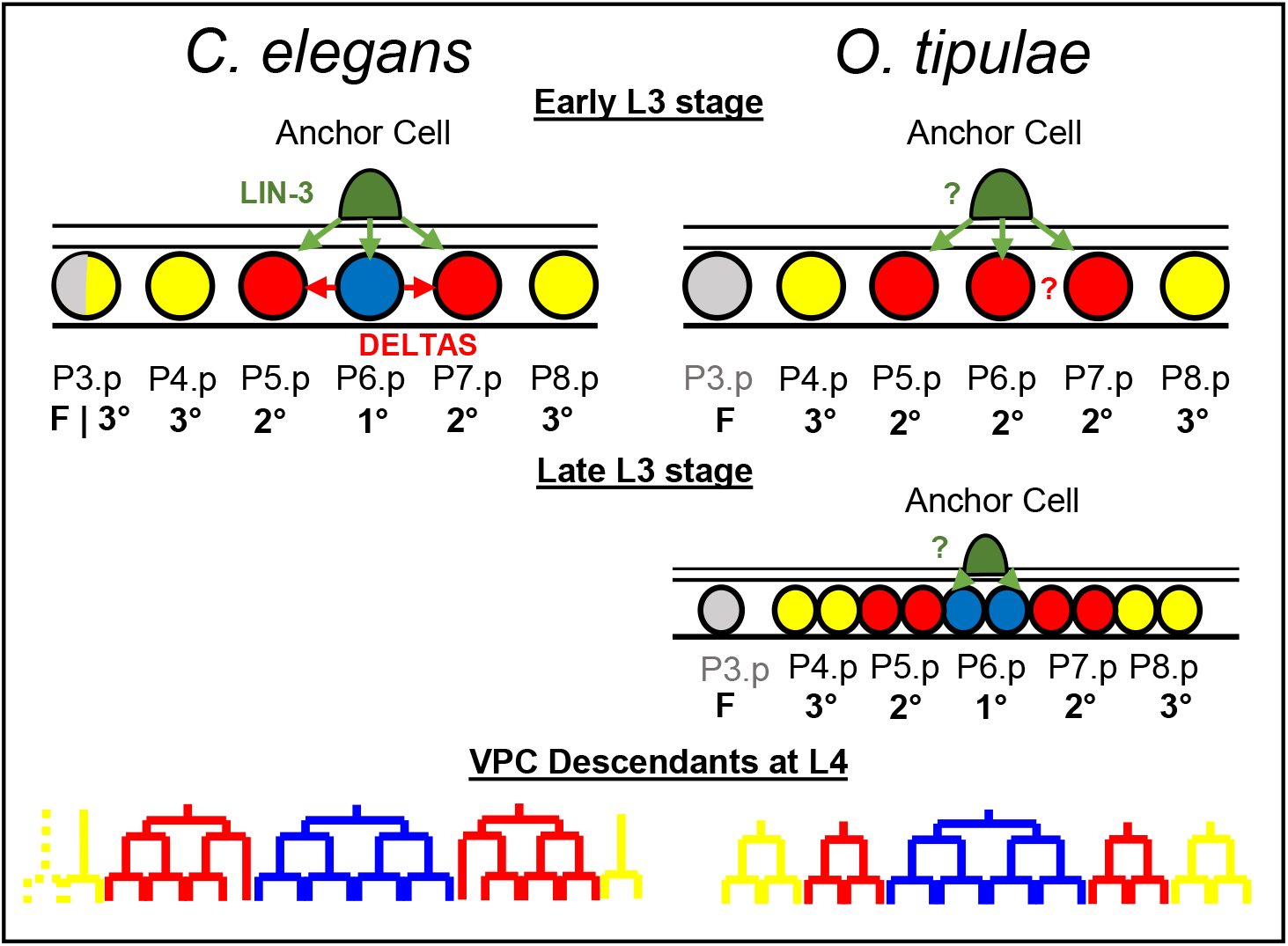
Vulval cell fate patterning in *Caenorhabditis elegans* and *Oscheius tipulae*. In the third larval stage (L3) of *C. elegans*, a cell from the somatic gonad known as the Anchor Cell (AC) produces an EGF-like inductive signal (LIN-3, green arrows) that activates the Ras pathway in the central vulva precursor cells (VPC). High Ras signaling promotes the 1° fate (blue circle) in P6.p which, in turn, produces Deltas (red arrows) which induce a 2° fate (red circle) and represses the 1° fate in P5.p and P7.p. Both fates prevent the formation of non-specialized epidermis (3° fate, yellow circles). Only VPCs with 1° or 2° fate will give rise to the cells that will form the vulva (bottom). P3.p is not competent to acquire a vulval cell fate (grey) in *O. tipulae*. Unlike in *C. elegans*, the AC of *O. tipulae* has been shown to be required after VPC division to induce the 1° fate in P6.p descendants. While a similar vulval cell fate pattern is conserved between the two nematodes, the cell division patterns of the 2° and 3° fates are different.

Among species with a two-step induction (Fig. 1), *Oscheius tipulae* is a rhabditid nematode found in the same habitat as *C. elegans* (Félix and Duveau 2012), which can be cultured in the same laboratory conditions (Félix et al. 2000b). A genetic screen was conducted ca. 20 years ago to isolate vulva development mutants in *O. tipulae* (Dichtel et al. 2001; Louvet-Vallée et al. 2003; Dichtel-Danjoy and Félix 2004b; Dichtel-Danjoy and Félix 2004a). This genetic screen led to a different spectrum of vulval cell fate and lineage phenotypes compared to those found in *C. elegans*. This result suggested a different sensitivity of the developmental system to mutation and therefore a different evolutionary potential. It also reflected the difference in development between *O. tipulae* and *C. elegans* (Dichtel-Danjoy and Félix 2004a). We then identified a null mutant in the Hox gene *lin-39*, with the same phenotype as in *C. elegans*, namely a loss of competence of the vulva precursor cells (Louvet-Vallée et al. 2003).

A draft of the *O. tipulae* genome has recently been published, along with a strategy to map the genomic location of loci whose mutation produces a visible phenotype (Besnard et al. 2017). As a proof of principle for the mutant identification technique, we described alleles of the *Oti-mig-13* locus with an unexpected vulva phenotype (Besnard et al. 2017). Here, we take advantage of the mapping approach to molecularly identify the collection of *O. tipulae* mutations affecting vulval cell fate patterning. We had found a single Induction Vulvaless locus with a single alelle and this turned out to be a *cis*-regulatory deletion in a tissue-specific enhancer of the *O. tipulae lin-3* homolog, which we confirmed by targeted CRISPR mutation of the element. We then identified mutations in Wnt pathway components *(mom-5/frizzled, mig-14/wingless*, and *egl-20/Wnt*) affecting fates of the *O. tipulae* vulva precursor cells, and discuss similarities and differences with *C. elegans* and *Pristionchus pacificus*, another nematode species where similar screens were conducted (Sommer 2006). Finally, the last class of vulval cell fate mutants caused an excess of 2°-fated cells. Unexpectedly, these mutations corresponded to lesions in *Oti-plx-1* and *Oti-smp-1*, encoding plexin and semaphorin, a cell signaling system that was not found in *C. elegans* vulva mutagenesis screens.

## Material and Methods

### Nematode culture

*C. elegans* and *O. tipulae* were handled according to usual procedures, on standard NGM plates with *Escherichia coli* strain OP50 as a food source (Brenner 1974; Félix et al. 2000b). *C. elegans* and *O. tipulae* strains were maintained respectively at 20°C and 23°C, unless otherwise indicated. N2 is used as a reference strain for *C. elegans* and CEW1, a wild isolate from Brazil, as a reference strain for *O. tipulae*. A list of strains used in this study is presented in Table S1.

### Mapping by sequencing and identification of molecular lesions

The mapping-by-sequencing strategy has been comprehensively described before (Besnard et al. 2017). In brief, each mutant *O. tipulae* line previously obtained in the CEW1 genetic background was crossed to males of the molecularly divergent wild isolate JU170. In the case of the fully Vulvaless *iov-1(mf86)* mutant, males of strain JU432 of genotype *iov-1(mf86); him(sy527)* were crossed to JU170 hermaphrodites. In all cases, individual F2 progeny with a recessive mutant phenotype were isolated and the mutant phenotype verified on the F3 brood. The pooled DNA of the progeny of mutant F2s was extracted using the Puregene Core Kit A (QIAGEN) and whole-genome sequenced at the BGI facilities. Pools from 37 to 152 individual F2s were used, depending on the ease of scoring of the mutant phenotype.

Sequencing reads from each mutant pool were mapped to the CEW1 genome using bwa (Li and Durbin 2009) and the resulting alignment converted to bam format using samtools (Li et al. 2009). Each mapping was further processed with the GATK suite (Van der Auwera et al. 2013) and allelic variants were called using HaplotypeCaller on a restricted list of JU170 sites for faster computation. Scaffolds having a mean JU170 allele frequency of less than 10% were selected as candidates for possibly linkage with a causative locus and processed for homozygous variant calling in an unrestrictive manner. JU170 variants were filtered out from the candidate scaffolds and the remaining variants were analyzed for any functional impact on the *O. tipulae* gene annotations (CEW1_nOt2) using snpEff (Cingolani et al. 2012). Scripts used for this processing pipeline can be found at: https://github.com/fabfabBesnard/AndalusianMapping. The candidate scaffolds were also analysed using Pindel (Ye et al. 2009) to identify large deletions or insertions, which were confirmed later by visual inspection with the Tablet software (Milne et al. 2013).

### Sanger sequence validation

The mutations identified by the mapping-by-sequencing approach were verified by Sanger sequencing of a PCR product. When other alleles of a given locus had been identified by genetic complementation screens, the gene was sequenced to find a possible lesion and in all cases we did find a lesion in the same gene. A list of primers used for sequencing can be found in Table S2.

### Identification of homologous genes

The predicted protein sequences of *O. tipulae* genes were obtained through the genome annotation (Besnard et al. 2017), now available from the Blaxter laboratory website: http://bang.bio.ed.ac.uk:4567. The sequence of their closest *C. elegans* homolog was identified using the BLASTP algorithm (Gish and States 1993), conditioning for highly similar alignments (>80% identity) and low e-value. Manual curation and re-annotation of the *O. tipulae* gene sequences were then performed using as a reference their closest *C. elegans* homolog. We aligned the amino-acid sequences of the re-annotated genes with their respective *C. elegans* homologs and outgroups using the Muscle algorithm implemented in MEGA X (Kumar et al. 2018) with default parameters. The phylogenetic relationship between the protein sequences was inferred using the Neighbor-Joining method (Saitou and Nei 1987) and tested for bootstrapping with 1000 replicates.

### Nomenclature

We followed *C. elegans* nomenclature and recommendations for other nematode species in Tuli et al. (2018). Briefly, mutant class names had been given at the time of our screen: *iov* for <u>i</u>nduction <u>o</u>f the <u>v</u>ulva; *dov*, for division of vulva precursor cells; *cov* for <u>c</u>ompetence and/or <u>c</u>entering of <u>v</u>ulva precursor cells. Once the molecular lesion has been identified, we use the name of the *C. elegans* homolog preceded by the species prefix for *Oscheius tipulae* ’Oti-’; for example the *iov-1(mf86)* allele is thus renamed *Oti-lin-3(mf86)*.

### Single molecule fluorescence *in situ* hybridization (smFISH)

smFISH in *O. tipulae* was performed as previously described (Barkoulas et al. 2016). Mixed-stage populations were used for mRNA localization experiment, while bleach-synchronized populations at the L3 larval stage were used for mRNA quantification. Only L3 stage nematodes with a gonad longer than 300 pixels (38.66 micrometers) were considered for mRNA quantification. The short fluorescently labelled oligos used in this study were acquired from LGC Biosearch Technologies and were used at a concentration of 100 to 200 mM. A list containing the sequences of the smFISH oligonucleotides is provided in Table S3.

### Phenotypic characterization and measurements of cell distances

The cell fates acquired by the *O. tipulae* vulva precursor cells were scored as previously (Dichtel et al. 2001). In summary, early L4 larvae were mounted with M9 solution on 4% agar pads containing 10 mM sodium azide and analyzed under Nomarski optics. Standard criteria were used to infer cell fates based on the topology and number of cells at different stages. Half fates were assigned when two daughters of the Pn.p cells acquired distinct fates after the first cell division.

Measurements of distances between the nuclei of Pn.p cells were performed on mounted larvae at 3 different developmental stages: L2 molt, early L3 (before the division of dorsal uterine DU cells), and mid L3 (after DU cell division and before Pn.p divisions). The distance between the center of the Pn.p and AC nuclei was measured in pixels using a Photometrics CoolSNAP ES camera and the Nikon NIS-Elements software (version 3.0.1). To avoid measurement errors due to the animal curvature, the distance between each Pn.p cell (except P6.p) and the AC was calculated via a Pythagorean formula. For example, the distance between P4.p and the AC is equal to:

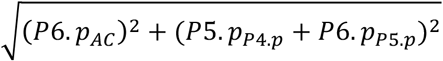

Where *P*6. *p_AC_* is the distance between P6.p and the AC, P5. and P4._p_, and *P*6.*p*_*P*5.*p*_ is the distance between P6.p and P5.p. Non-normalized measurements can be found in Table S4.

### Genome editing

We followed the CRISPR-Cas9 target design in Paix et al. (2015). We targeted the following sequence at the *O. tipulae lin-3* cis-regulatory region 5’-<u>cCACCTG</u>catgtcctttttgcgc-3’ (E-box site in uppercase, within an underlined NGGNGG PAM motif in the negative strand). The *mf113* allele was produced with the synthetic Oti_lin-3_A-2 -GCGCAAAAAGGACAUGCAGG-crRNA manufactured by Dharmacon (GE Healthcare), while *mf114* was produced with the same crRNA sequence synthetized by IDT. Each crRNA was mixed with tcRNA (Paix et al. 2015) at an equimolar concentration of 200 micromoles/microliter. The tcRNA:crRNA mix was incubated in a thermal ramp between 95 and 25°C, decreasing by 5°C every two minutes, and then mixed with purified CRISPR-Cas9 protein in HEPES buffer (pH 7.4), reaching a final concentration of 30 μM of the tcRNA:crRNA duplex and ~18 μ.M of purified protein. The final mix was incubated for 15 minutes at 37°C and then injected into the gonad of *O. tipulae* gravid adults. The F1 progeny of the injected nematodes were placed into new plates and, after letting them lay eggs for one day, screened for deletions by PCR with the mf86-EboxA-F and mf86-R primers. Heterozygous F1 animals were identified by band-size separation on 3% agarose gels, and homozygous F2 mutants were easily spotted by their bag phenotype. Only a single mutation per injection session (> 10 P0s and > 200 F1s) was obtained.

### Immunofluorescence staining

Bleach-synchronized larvae and mixed-stage populations were fixed and permeabilized for immunostaining using previously described methods (Louvet-Vallée et al. 2003; Kolotuev and Podbilewicz 2004; Kolotuev and Podbilewicz 2008). In brief, OP50-grown populations were washed 3 times in distilled water and placed onto poly-L-lysine-coated (SIGMA P0425-72EA) slides prior to freeze-cracking. Worms with an open cuticle were incubated in antibody buffer with the mouse MH27 antibody against the epithelial cell adherent junctions (Francis and Waterston 1991). This antibody was obtained from the DHSB and used at a concentration of 1 mg/mL. As secondary antibody, we used the goat anti-mouse antibody from Abcam labelled fluorescently with Alexa Fluor 488 (ref. #ab150113). The slides containing immunofluorescently labelled worms were mounted with GLOX buffer (Ji and van Oudenaarden 2012) containing DAPI, covered with a cover slip, and imaged with a PIXIS camera (Princeton Instruments).

### Data and reagent availability

Supplementary Tables are available throguh the FigShare portal:

- Table S1. List of strains used in this study.
- Table S2. Sequences of DNA primers used in this study. Sequencing primers to verify by Sanger sequencing the mutations identified by the mapping by sequencing approach, and to identify the molecular lesion in additional alleles.
- Table S3. Sequences of smFISH probes used in this study. The fluorophore coupled to each probe is noted at the end of the set name.
- Table S4. smFISH quantifications, distance measurements and vulval cell fates used in this study.

Data and strains are available by contacting Marie-Anne Félix. Code for mutant identification is available at https://github.com/fabfabBesnard/Andalusian_Mapping.

## Results

### The sole hypoinduction mutation is due to a cis-regulatory change in *Oti-lin-3*

Our prior mutagenesis screens had yielded a single mutant with an Induction Vulvaless phenotype, i.e. the 1° and 2° fates are transformed to a 3° fate (two rounds of division and fusion to the hyp7 syncytium, represented in yellow in the figures) but rarely to a non-competent state (fusion to hyp7 without division, prior to the L3 stage, represented in grey) (Dichtel-Danjoy and Félix 2004b). This allele, *iov-1(mf86)*, was obtained after TMP-UV (trimethylpsoralene-ultraviolet) mutagenesis. The mapping-by-sequencing approach identified a 191 bp deletion upstream of the coding sequence (second ATG) of *O. tipulae lin-3 (Oti-lin-3)* (Fig. 2B). We hypothesized that this deletion may cause a reduced level of expression in *Oti-lin-3* and thus performed single molecule Fluorescent In Situ Hybridization (smFISH) experiments to quantify *Oti-lin-3* mRNA number (Raj et al. 2008; Barkoulas et al. 2013; Barkoulas et al. 2016). Indeed, the *Oti-lin-3* mRNA level in the anchor cell was much decreased in animals bearing the *Oti-lin-3(mf86)* deletion compared to animals of the CEW1 reference strain (Fig. 2C, Kolmogorov-Smirnov test, p<10^-9^). The deleted region in *Oti-lin-3(mf86)* contains an E-box motif known to be conserved in *Caenorhabditis* species (Fig. 2B) (Barkoulas et al. 2016), as well as a second less characteristic putative E-box motif.

**Figure 2.**
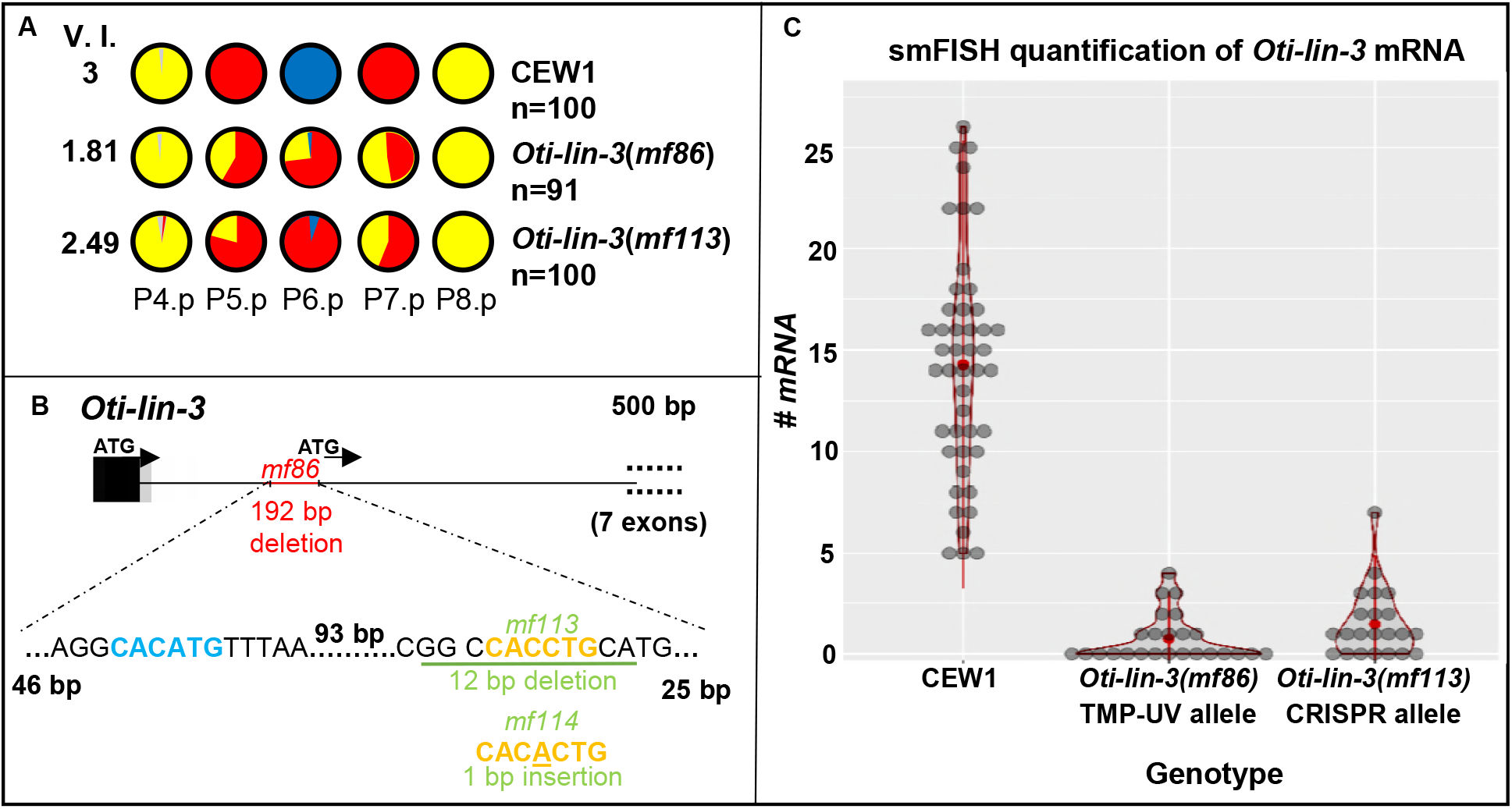
Cis-regulatory lesions in *Oti-lin-3/EGF* cause a hypo-induction of 1° and 2° vulval cell fates. (A) P(4-8) cell fates in the wild-type CEW1 *O. tipulae* reference strain and mutants for *lin-3/EGF*. The pie diagrams represent the percentage of cell fates over individuals. Yellow, red, and blue are for the 3°, 2° and 1° fates, respectively. Grey denotes an undivided cell fused to the hypodermis. The Vulva Index (V.I.) is calculated as the average number of cells acquiring a vulval cell fate in a set of animals. The quantifications of *Oti-lin-3(mf86)* are from (Dichtel-Danjoy and Félix 2004b). (B) Position of the deletions in the TMP-UV and CRISPR alleles. As in *C. elegans*, the *lin-3* gene of *O. tipulae* is predicted to have two alternative ATGs, with the anchor cell cis-regulatory element upstream of the second ATG. Note that the seven exons following the second ATG were excluded from the diagram. (C) Distributions of *Oti-lin-3* mRNA number in the anchor cell of wild-type CEW1 and *lin-3* cis-regulatory mutants, as quantified by smFISH.

To test whether the conserved E-box motif was required for the expression of *Oti-lin-3* and also confirm that the 191 bp deletion was causal for the vulva phenotype, we performed a CRISPR/Cas9 experiment specifically targeting this site. We obtained two new mutations, a smaller 12 bp deletion (*mf113*) and a one-bp insertion in the E-box (*mf114*). Both showed a strong decrease in the level of induction, confirming that the lesion in the *Oti-lin-3* gene is causal for the phenotype (Fig. 2A, Table S4). Further smFISH experiments in the *Oti-lin-3(mf113)* mutant revealed a similar level of mRNAs as in the *Oti-lin-3(mf86)* mutant (Kolmogorov-Smirnov test non-significant, *p*=0.94). We conclude that the conserved E-box site is also required in *O. tipulae* for *lin-3* expression and that LIN-3 secreted from the anchor cell is necessary for induction of both 2° and 1° fates.

### The Wnt pathway plays a role in vulva precursor competence/induction and fate pattern centering

A large class of mutants in our screen displayed a lower number of competent Pn.p cells (transformation to 4°/grey fate) and a displacement of the 1° fate from P6.p to P5.p. In *C. elegans*, this phenotype has not been seen at this high level of penetrance. The mapping-by-sequencing approach had already identified one locus in this class as *Oti-mig-13* (Besnard et al. 2017). We further identified in this class mutations in two Wnt pathway components:

I. i) a Wnt receptor gene, *Oti-mom-5* (supported by two alleles, including an early stop) (Fig. 3B). Relationships among Wnt receptors paralogs in the different species is shown in Fig. S4. Curiously, the *Oti-mom-5* putative null allele, *sy465*, is not embryonic lethal in *O. tipulae*, while it is lethal in *C. elegans* (embryonic mesoderm versus endoderm specification; Rocheleau et al. 1997).
II. ii) a Wnt processing protein, *Oti-mig-14* (homolog of *Drosophila* Wntless) (Bänziger et al. 2006; Yang et al. 2008). The *mf34* allele is an amino-acid substitution and likely a hypomorph that may negatively affect the activity of all Wnts.

**Figure 3.**
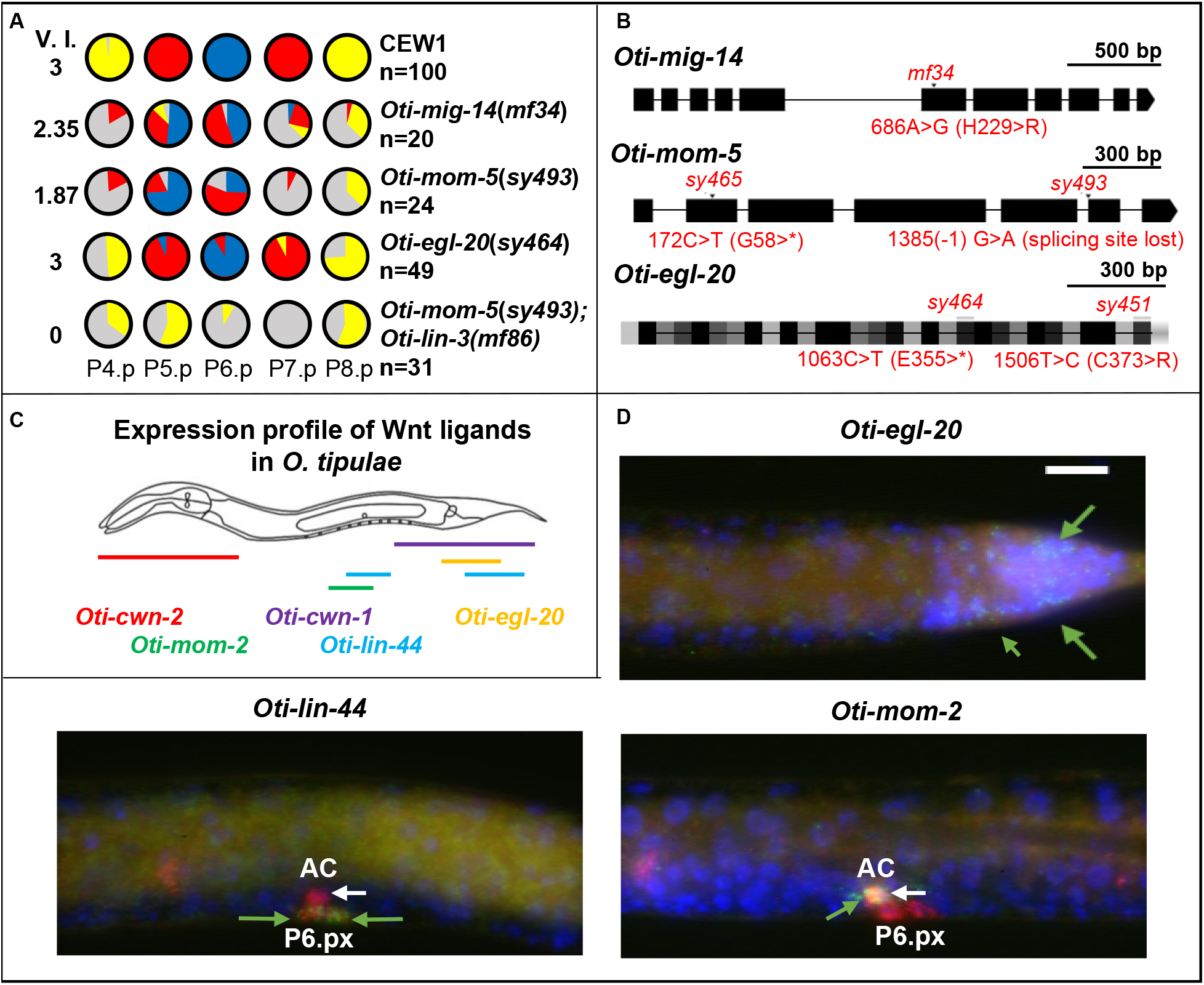
*O. tipulae* mutants in Wnt signaling display defects in competence and centering of the 1° fate on P5.p. (A) Pie diagrams representing the percentage of P(4-8).p cells acquiring one of the four possible cell fates (blue, red, yellow and grey for the 1°, 2°, 3° or 4° fates, respectively) for animals of different genotypes. The Vulva Index (V.I.) is calculated as the average number of cells acquiring a vulval cell fate in a set of animals. The quantifications in *Oti-mig-14(mf34)* and *Oti-mom-5(sy493)* animals are from (Louvet-Vallée et al. 2003) and that of *Oti-egl-20(sy464)* from (Dichtel et al. 2001). (B) Position of different mutations on genes encoding Wnt pathway components. A star designates a stop codon. (C) Diagram of Wnt ligand expression profiles in *O. tipulae* at mid L3 stage. smFISH images of *Oti-cwn-1* and *Oti-cwn-2* are found in Fig. S3. D) smFISH images of *Oti-egl-20, Oti-lin-44* and *Oti-mom-2* Wnt ligands after P6.p division at the L3 stage. mRNAs are visible as green dots. The animals were also labeled with DAPI (in blue, labeling nuclei) and fluorophore probes for *Oti-lag-2/delta* (in red, labeling the anchor cell, P6.p descendants, and distal tip cells outside the field of view). *Oti-egl-20* is visible only at the posterior part of the animal (green arrows). *Oti-mom-2* mRNAs (green arrow) are found in the anchor cell (white arrow), while *Oti-lin-44* mRNAs (green arrows) appear in P6.p daughters (as well as sex myoblast precursors outside the focal plane). All the images are set to the same scale. The size of the white bar is 10 micrometers. Anterior is to the left in all images, and the ventral side down.

We had distinguished somewhat arbitrarily classes of vulva mutations that affect competence and centering (cov mutants) from those that affect division of vulval precursor cells *(dov* mutants) (Dichtel et al. 2001). Among the latter class, we found that the *dov-4* locus encodes a Wnt-type ligand, *Oti-egl-20* (supported by two alleles, including a premature stop). The *Oti-egl-20* mutation results in a lower competence and division frequency of P4.p and P8.p, but hardly affects P(5-7).p. Centering of the 1° fate on P5.p only occurs at low penetrance. Overall, the *Oti-egl-20* phenotype is similar to that of *Oti-mig-14* or *Oti-mom-5*, albeit much weaker, suggesting the involvement of other Wnt family ligands.

The *O. tipulae* genome encodes five genes coding for Wnt signaling molecules, which we found to be 1:1 orthologs to the five Wnt genes in *C. elegans* (Fig. S4). By smFISH, the expression pattern of each of these five genes was found to be quite similar in L1-L3 larvae to that of each ortholog in *C. elegans*, as determined in Song et al. (2010) and Harterink et al. (2011). Specifically, *Oti-egl-20* is expressed in the posterior region of the animal from the L1 stage (Fig. 3D). *Oti-cwn-1* is also expressed quite posteriorly (Fig. S3A). *Oti-cwn-2* is expressed in the anterior region (Fig. S3B). *Oti-mom-2* is expressed in the anchor cell from the L3 stage (Fig. 3D). *Oti-lin-44* is expressed in the tail region and, in the L3 stage, in P6.p daughters (Fig. 3, Fig. S6). Similar to *cwn-1* in *C. elegans* (Harterink et al. 2011; Minor et al. 2013), we found that *Oti-lin-44* is in addition expressed in the sex myoblast precursors that are located left and right of the anchor cell in the L3 stage (Fig. S6). As the sex myoblast expression of *Oti-lin-44* differed from the reported uterus/anchor cell pattern in *C. elegans* using lacZ staining or fluorescent reporters (Inoue et al. 2004), we localized *lin-44* by smFISH in *C. elegans* and saw a similar expression in the sex myoblasts (identified by labeling with *hlh-8::GFP;* Harfe et al. 1998) and P6.px, and none in the uterus and anchor cell (Fig. S7). In conclusion, the larval expression patterns of the five Wnt genes were thus similar in *O. tipulae* and *C. elegans*.

From the *Oti-egl-20* expression pattern and mutant phenotype, the EGL-20 protein is produced from the posterior of the animal and promotes Pn.p cell competence as far as P4.p. P3.p is not competent and does not divide in *O. tipulae* (Félix and Sternberg 1997; Delattre and Félix 2001) and is thus not affected by Wnt pathway mutations, whereas it is highly sensitive to Wnt pathway modulation in *C. elegans* (Pénigault and Félix 2011b). However, the difference in phenotype severity between *Oti-mig-14* or *Oti-mom-5* mutants on one hand and *Oti-egl-20* (including the *sy464* allele with a stop codon) on the other hand, suggests that other Wnt signals, perhaps mostly CWN-1 from the posterior as in *C. elegans* (Gleason et al. 2006), may act jointly to promote Pn.p competence.

Overall, the major differences between *C. elegans* and *O. tipulae* for this class of mutants are 1) Wnt pathway mutations were not found in the first vulva mutant screens in *C. elegans;* 2) the miscentering of the 1° fate on P5.p is much more penetrant in *O. tipulae* than in *C. elegans* (Fig. 3, see Discussion). 3) Wnt pathway mutations lead to low division frequency of P8.p in *O. tipulae* compared to *C. elegans*, for a comparable or even weaker effect on P4.p: *Oti-egl-20(sy464)* and *Cel-egl-20(n585)* animals show 30% and 1% loss of division of P8.p, respectively (Dichtel et al. 2001; Myers and Greenwald 2007).

### The hyperinduced mutations affect plexin and semaphorin genes

Much more unexpected is the identification of the mutations resulting in a vulva hyperinduction phenotype. Indeed, the *iov-3* locus turned out to correspond to the *Oti-plx-1* gene, coding for a plexin (one small deletion and two missense alleles), while the *iov-2* mutant shows a deletion in the *Oti-smp-1* gene, coding for a semaphorin-type ligand (Fig. 4A). This implicates a new intercellular signaling pathway in vulval cell fate patterning and induction.

**Figure 4.**
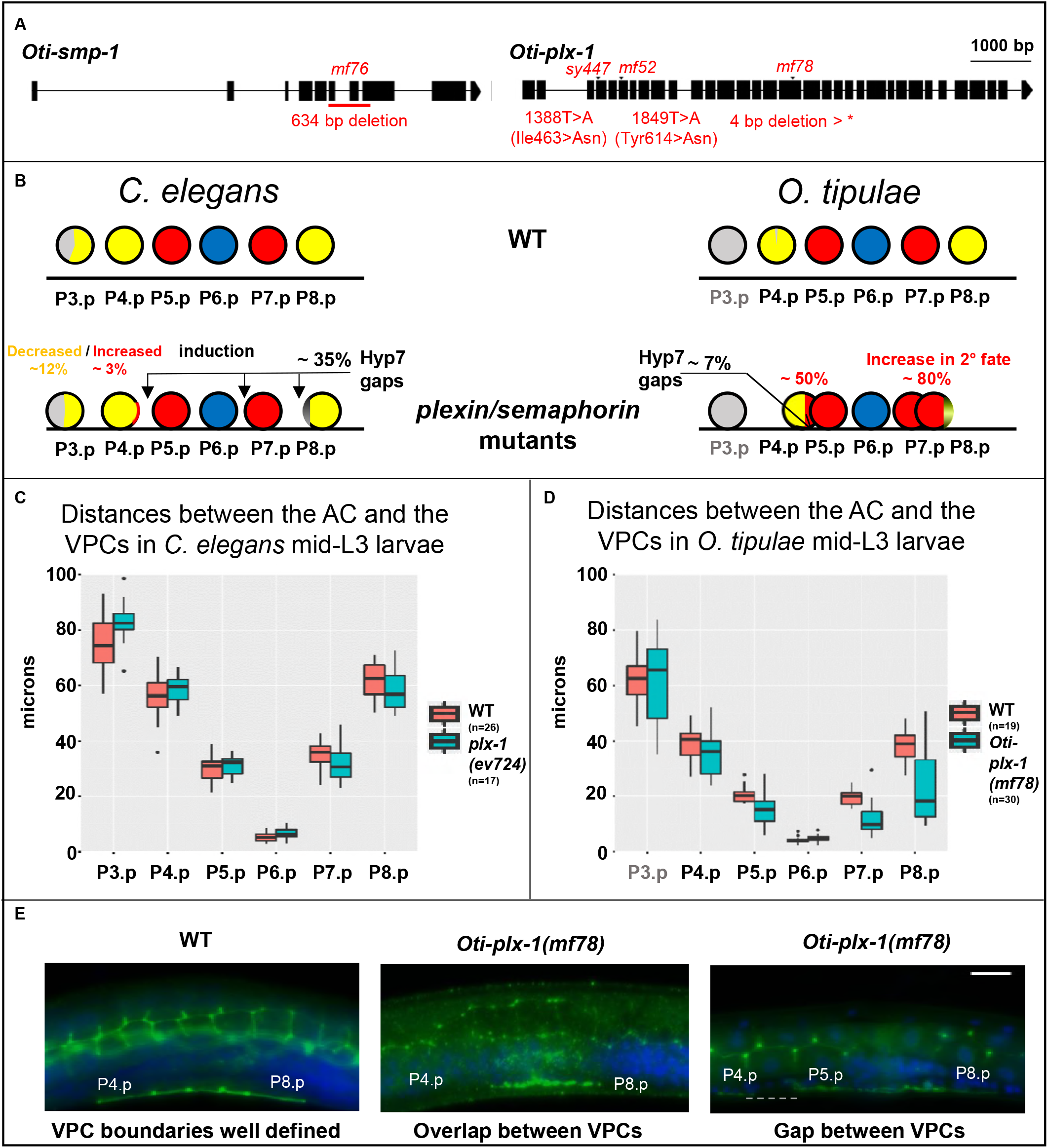
*O. tipulae plexin/semaphorin* mutants present defects in vulva induction and closer VPC cells. (A) Gene models of *Oti-smp-1* and *Oti-plx-1* with their respective mutations in *O. tipulae*. (B) Schematic depiction of the phenotypic effects of plexin/semaphorin mutants in *C. elegans* and *O. tipulae* on the induction and the localization of Pn.p cells. Quantifications can be found in Table S4. Arrows show the most common localization of intercellular space (gaps) between vulva precursor cells. Each vulva precursor cell diagram (circle) is colored according to the frequency of its acquired fate (yellow, red, and blue for the 3°, 2° and 1° fates, respectively, and grey for undivided). Data from Dichtel-Danjoy & Félix 2004. (C) Normalized distances between the AC and the VPCs in *C. elegans* wild type and *plx-1(ev724)* animals at the mid-L3 stage after DU division. Only the distances between the AC and P3.p, and P6.p are significantly larger in *plx-1* mutants compared to wild-type (Wilcoxon rank sum test, *p* < 0.05). (D) Normalized distances between the AC and the VPCs in *O. tipulae* wild type and *Oti-plx-1(mf78)* animals at the mid-L3 stage after DU division. Distances between each of P(4-8).p and the AC, except for P6.p, are all significantly smaller in *Oti-plx-1(mf78)* mutants relative to wild-type (Wilcoxon rank sum test, with p-values < 10^-5^). (E) Immunostaining of cell junctions with MH27 antibody (in green), with DAPI staining in blue. The central panel shows overlapping VPCs, while the right panel shows a rare instance of a gap (dotted line) in *Oti-plx-1(mf78)* animals. All the images are set to the same scale. The size of the white bar is 10 micrometers. Anterior is to the left in all images, and the ventral side down.

The plexin-semaphorin pathway is well known for contact-dependent growth inhibition between neurons, acting in many organisms (Kolodkin et al. 1992; Luo et al. 1993; Winberg et al. 1998). In *C. elegans*, mutations in *smp-2/mab-20, smp-1* and *plx-1* (Roy et al. 2000; Ginzburg et al. 2002; Fujii et al. 2002; Dalpé et al. 2004; Pickett et al. 2007; Nukazuka et al. 2008) were found and mostly studied for their effect on the displacement of sensory organs (rays) in the male tail. Their impact on vulva formation mostly concerns late morphogenesis events that take place after the three rounds of Pn.p divisions (Liu et al. 2005; Dalpé et al. 2005; Pellegrino et al. 2011), while their effect on vulval induction is minor (Liu et al. 2005), as also shown in Fig. 4B and Table S4.

The hyperinduction of P4.p and P8.p in the *O. tipulae iov-2/smp-1* and *iov-3/plx-1* mutants is a transfomation of 3° to 2° fate. The ectopically induced cells never adopt a 1° fate; they join the main vulval invagination and therefore the adult phenotype is a protruding vulva and not additional bumps on the cuticle as in the *C. elegans* Multivulva mutants. This contrast with the *C. elegans* hyperinduced mutants, which correspond to an excess of Ras pathway signaling, leading to ectopic 1° and 2° fates.

To understand why plexin and semaphorin mutations cause a vulval hyperinduction in *O. tipulae*, we measured cell position at the time of induction, before the formation of the vulval invagination. As in *C. elegans* plexin and semaphorin mutants, we observed that the vulva precursor cells do not form an antero-posterior row as in wild-type animals (Liu et al. 2005; Dalpé et al. 2005) but instead either overlap left and right of each other or sometimes show a lack of junction and a gap between successive cells (Fig. 4B,C,D and E). In contrast to *C. elegans*, gaps are rare in *O. tipulae* mutants and do not concern the three central cells. Instead, left-right overlaps occur between P4.p and P5.p, and between P7.p and P8.p. As these overlaps could alter the distance between the anchor cell and the Pn.p cells, we measured these distances and found that they were shorter in the *O. tipulae plx-1(mf78)* mutant but not in the *C. elegans* counterpart *plx-1(ev724)* (Fig. 4C, D). (Both alleles are deletion alleles, and thus putatively comparable null alleles.) As a consequence, the vulva precursor cells tend to be closer to the anchor cell in *O. tipulae*, liklely explaining the excess of 2° fate induction in the first induction wave.

In summary, the identification of these four different mutations points to plexin/semaphorin signalling as an important pathway for the correct induction of the vulva precursor cells, due to its effect on vulval precursor cell positioning.

## Discussion

### The unsurprising single Vulvaless mutation in *O. tipulae*

In the first *C. elegans* screens for vulval induction defects, most Vulvaless mutations corresponding to induction defects affected the genes *lin-2, lin-7* or *lin-10* (Horvitz and Sulston 1980; Ferguson and Horvitz 1985; Ferguson et al. 1987). Only rare tissue-specific reduction-of-function alleles were recovered in *lin-3* and *let-23*, coding for the EGF and the EGF receptor, respectively. Downstream factors in the EGFR-Ras/MAP kinase cascade were only subsequently obtained by suppressor or enhancer screens (Sternberg and Han 1998).

For *lin-3*, the first *C. elegans* allele, *e1417*, turned out to be a base substitution affecting a cis-regulatory E-box (Hwang and Sternberg 2004). The second viable allele, *n378*, is a substitution in the signal peptide, showing high tissue-specificity for reasons still ignored (Liu et al. 1999). Further *lin-3* alleles were obtained in non-complementation screens or screens for lethal mutants (Ferguson and Horvitz 1985; Liu et al. 1999). In summary, besides the *lin-2/lin-7/lin-10* genes, a main target for a Vulvaless mutation appeared to be the cis-regulatory element that activates *lin-3* expression in the anchor cell in a tissue-specific manner. Given this, the sole Vulvaless mutation we found in mutagenesis of *O. tipulae, iov-1(mf86)*, is a remarkably predictable hit: a deletion in a homologous non-coding region to that mutated in *Cel-lin-3(e1417)* (Barkoulas et al. 2016). Random mutagenesis ended up being as targeted as the CRISPR/Cas9 experiment that confirmed the importance of this E-box (Fig. 2).

Concerning *lin-2, lin-7* or *lin-10*, we now know that the proteins LIN-2/CASK, LIN-7/Velis and LIN-1/Mint1 bind to the C-terminus of the LET-23/EGFR receptor and help to localize it to the basolateral membrane facing the anchor cell (Simske et al. 1996; Kaech et al. 1998). Mutations in any of these three loci were so far not recovered in *C. briggsae* and *P. pacificus* nor here in *O. tipulae* (Fig. 5). It is thus likely that either their loss of function is lethal or it does not affect the vulva. It will be interesting to delete them using reverse genetic methods such as CRISPR/Cas9 mediated genome modification.

**Figure 5.**
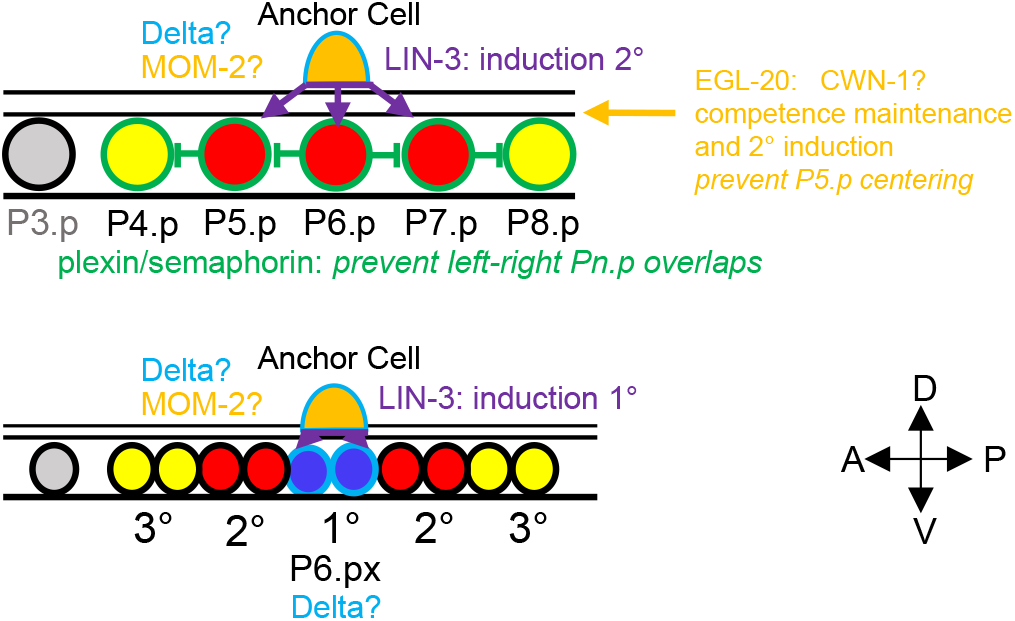
Expression of signaling molecules and vulval cell fate patterning in *Oscheius tipulae*. The VPCs are color-coded according to their fate as in previous figures. Their boundary is color-coded according to the signaling molecules that they express (at least as mRNAs): LIN-3 in purple, Wnts in orange, plexin in green and Delta in light blue. A question mark indicates that the effect of removing this signal is not known. Note that in additionthe sex myoblasts left and right of the AC express *lin-44/Wnt*.

### A surprise signaling pathway found only in *O. tipulae* vulva mutant screens

In stark contrast, the identification of the semaphorin-plexin pathway using the hyperinduced mutations in *O. tipulae* was unpredictable and is a novel result. This genetic screen outcome could not have been foreseen from results in *C. elegans, C. briggsae* (Seetharaman et al. 2010; Sharanya et al. 2015; Sharanya et al. 2012) nor *P. pacificus* (Jungblut and Sommer 1998; Jungblut and Sommer 2001; Schlager et al. 2006; Tian et al. 2008). In the case of *C. elegans*, the vulval fate specification errors in plexin/semaphorin mutants are indeed rare and occur at low penetrance and in directions of both excess and loss of induction. Instead in *O. tipulae*, the specification of P4.p or P8.p as a 2° fate is quite penetrant and we only observe hyperinduction (Table S4, Fig. 4B). The cell positioning defects in the *O. tipulae* plexin/semaphorin mutants explain that the hyperinduction of vulval fates is gonad-dependent (Dichtel-Danjoy and Félix 2004b). In contrast, in *C. elegans* hyperinduced mutants, such as *lin-1, lin-13, lin-15, lin-31* and *lin-34(d)*, retain some vulval induction upon anchor-cell ablation (or in *lin-3* double mutants) (Ferguson et al. 1987; Han and Sternberg 1990).

What explains the difference between *C. elegans* and *O. tipulae* in the effect of mutations in the plexin-semaphorin pathway? In both species, semaphorin and plexin appear to act in contact inhibition of the VPCs while they grow and contact each other (Liu et al. 2005) (Fig. 4). We propose that two not mutually exclusive phenomena concur to the fate specification difference. First, the VPCs are in average closer to the anchor cell in the early L3 stage in *Oti-plx-1* mutants compared to the corresponding *C. elegans plx-1* mutants (Fig. 4C,D); this likely increases the exposure of P4.p and P8.p to Oti-LIN-3 from the anchor cell, hence the 2° fate. The smaller body size of *O. tipulae* may also play a role. Second, the 2° fate is in part induced in *C. elegans* by direct contact between P6.p and other VPCs through transmembrane Delta ligands. In *O. tipulae*, due to the difference in fate patterning mechanism, we have no evidence of lateral signaling, whereby the 1°-fated cell induces the 2° fate in its neighbors, nor of Notch pathway involvement, except maybe later through *Oti-delta* expression in P6.p daughters; indeed, P5.p, P6.p and P7.p do not appear different from each other before their division – although this may be due to the lack of adequate markers (Félix and Sternberg 1997). Signaling from the anchor cell at a distance is thus potentially stronger in *O. tipulae* than in *C. elegans*.

In *C. elegans*, vulva precursor cells are attracted towards the anchor cell in response to LIN-3 signaling, thus creating a positive feedback whereby the most induced cell moves closest to the anchor cell (Grimbert et al. 2016). The same feedback may be at stake for the 2° cells, but curiously, we never observed an excess of 1°-fated cells in *O. tipulae*. This correlates with the fact that we do not observe other VPCs overlapping with P6.p nor contacting the anchor cell in the plexin/semaphorin mutants. It is thus possible that a lateral inhibition from P6.p to its neighbors takes place in these mutants, preventing the positioning of two VPCs below the anchor cell.

### Wnt and EGF pathways act jointly in vulval competence and induction

We find that *O. tipulae* Wnt pathway mutants affect Pn.p competence and induction (2° to 3° and 3° to F transformations, Fig. 3A) and result in centering of the 1° fate on P5.p. The initial genetic screens for *C. elegans* vulva mutants did not identify the Wnt pathway. The corresponding mutants were found later by specifically screening for mutants that had a variably expressed protruding vulva phenotype (Eisenmann et al. 1998; Eisenmann and Kim 2000). It would be tempting to conclude at a difference in Wnt pathway involvement in *O. tipulae* compared to *C. elegans* vulva induction. However, we propose that the difference is subtle.

In *C. elegans*, the Wnt pathway is mostly known to maintain vulval precursor competence to receive the LIN-3 signal in the L2 and L3 stage (Eisenmann et al. 1998). In the absence of Wnts, the Pn.p cells adopt a F fate (Fusion with hyp7 in the L2 stage) instead of the 3° fate (one division in the L3 stage before fusion to hyp7) (Gleason et al. 2006). This prevents them from being induced to a vulval fate. In other words, the Wnt signaling pathway establishes competence (F to 3° fate transformation) for the next round of signaling (EGF, which induces 1° and 2° fates). Yet the two inductions by Wnt and EGF in *C. elegans* are partially intermingled. Indeed, the Wnt pathway also participates to the induction of 2° vulval fates versus the 3° fate (Eisenmann et al. 1998; Gleason et al. 2002; Seetharaman et al. 2010; Milloz et al. 2008; Braendle and Félix 2008). Conversely, the LIN-3/EGF pathway participates to the "competence maintenance" (F versus 3°) (Myers and Greenwald 2007). Thus, both pathways appear to jointly act in *C. elegans* to promote both “competence” (a very first induction) and 2° vulval fate induction.

The same holds true in *O. tipulae*, with quantitative variations in mutant phenotypes. In the *Oti-lin-3(mf86)* mutant, the 1° fate is abolished while the 2° fate is reduced. The intermediate level of 2° fate may be due to some remaining *Oti-lin-3* gene expression (Fig. 2C). Alternativey, another signal, such as Wnts, may participate to 2° fate induction. Accordingly, a double mutant between EGF and Wnt pathways, *Oti-mom-5(sy493); lin-3(mf86)*, abolishes induction, as in *C. elegans* (Eisenmann et al. 1998; Braendle and Félix 2008) (Fig. 3A). We thus conclude that despite quantitative differences in mutant penetrance, the joint involvement of the Wnt and EGF pathways in the induction of vulval fates appears similar in *C. elegans* and *O. tipulae*.

This joint induction by Wnts and LIN-3 differs from the situation described in an outgroup nematode, *Pristionchus pacificus* (Kiontke et al. 2007). In this species, the induction of vulval fates occurs gradually before and after Pn.p divisions (2° then 1°), as in *O. tipulae* and unlike *C. elegans* (Sigrist and Sommer 1999; Kiontke et al. 2007). There is no equivalent to the 3° fate in *P. pacificus*. Indeed, on the anterior side non-competent cells die by apoptosis. On the posterior side, P8.p is competent early on to replace P(5-7).p then fuses to hyp7 without division – only after the onset of vulval induction, which occurs earlier than in *C. elegans* compared to larval molts (Sommer 1997; Sigrist and Sommer 1999; Jungblut and Sommer 2000). Only two β-catenins were found in *P. pacificus* (Tian et al. 2008) (there are no *wrm-1* or *sys-1* orthologs). The *Ppa-bar-1/armadillo(0)* mutant obtained by a targeted reverse genetic approach is maternal-effect lethal (unlike in *C. elegans)* and strongly affects the level of induction (Tian et al. 2008). As in *C. elegans*, the multiple Wnt-ligands and receptors are partially redundant (Gleason et al. 2006; Tian et al. 2008). The Ppa-LIN-44 protein is said from alkaline phosphatase reaction to be expressed in the uterus (Tian et al. 2008). It may be good to clarify whether this expression may be in the sex myoblasts on either side of the uterus, as observed in *O. tipulae* and *C. elegans* (Fig. 3, S6). This is important, as Ppa-LIN-44 cannot represent the vulva induction signal as proposed if it is not expressed in the gonad precursors ablated in Sigrist and Sommer (1999). Indeed, the only other Wnt expressed in the *P. pacificus* gonad is *Ppa-mom-2*, but its expression in the anchor cell appears to start much after the induction of 2° fates begins (Sigrist and Sommer 1999; Kiontke et al. 2007; Tian et al. 2008).

### The Wnt pathway is required for correct centering of the vulval pattern

The clearest difference of Wnt pathway phenotypes between *C. elegans* and *O. tipulae* lies in the centering of the 1° fate on P5.p, and the likely correlated higher penetrance of the F fate in P7.p. In *C. elegans*, only a small percentage of Wnt pathway mutant animals displays P5.p centering, which was shown to reflect the posterior displacement of P6.p compared to the anchor cell, and a higher variance in cell positions (Milloz et al. 2008; Grimbert et al. 2016). In *Oti-mom-5* animals, a strong shift in anchor cell position relative to P6.p and P5.p in the L2 stage was also observed (Louvet-Vallée et al. 2003). Quantitative differences between the various phenotypes in the two species likely correspond to the extent of cell displacement.

## Conclusions

We present in Fig. 5 our current model of the vulval cell fate patterning mechanism in *O. tipulae*. Oti-LIN-3 produced by the anchor cell is important for induction of 1° and 2° fates. Oti-LIN-3 is thus likely the inductive signal for both steps of induction as defined in Félix and Sternberg (1997). The 1° fate induction appears to always occur upon contact with the anchor cell, which may represent the requirement for a transmembrane ligand or simply high concentration of the ligand.

The Wnt pathway is required for the F to 3° induction and also for the 3° to 2° induction (directly or indirectly), as in *C. elegans*. Wnts also prevent centering of the vulva pattern on P6.p, probably by a repulsive action of the posterior Wnts (Fig. 6). The latter is much more evident in *O. tipulae* than in *C. elegans* (Grimbert et al. 2016).

Our findings on the effects of both Wnt and semaphorin pathways on VPC positioning relative to the anchor cell emphasize the importance of cell positioning in vulval cell fate patterning since gradients of signaling molecules (EGF, Wnt) are involved. We note that the Wnt pathway mutants and *Oti-mig-13* have similar vulva phenotypes, as is true for their effect on Q_r_ neuroblast migration (Sym et al. 1999; Wang et al. 2013). The VPC positioning defect may link these regulatory pathways to cell polarity, growth and movement, and to the actin cytoskeleton (Wang et al. 2013; Grimbert et al. 2016).

**Table 1.** *Oscheius tipulae* vulva loci identified by mapping-by-sequencing approach.

| Locus | Allele | Phenotypes | Mutagen | Position* | Mutation | Oti gene | Cel homolog | Type of lesion | Reported before in |
| --- | --- | --- | --- | --- | --- | --- | --- | --- | --- |
| *cov-4* | *sy465* | Competence loss,<br>P5.p centering | EMS | 10:202548 | G/A | g06014 | *mom-5* / *frizzled* | Premature stop | Louvet-Vallée et al. 2003 |
| *cov-4* | *sy493* | Competence loss,<br>P5.p centering | EMS | 10:201110 | C/T | g06014 | *mom-5* / *frizzled* | Splice acceptor | Louvet-Vallée et al. 2003 |
| *cov-5* | *mf34* | Competence loss,<br>P5.p centering | EMS | 3:189504 | T/C | g01986 | *mig-14* / *Wntless* | Missense variant | Louvet-Vallée et al. 2003 |
| *dov-4* | *sy464* | P4.p/P8.p do not divide,<br>some P5.p centering | EMS | 4:882344 | G/A | g02936 | *egl-20* / *Wnt* | Premature stop | Dichtel et al. 2001 |
| *dov-4* | *sy451* | P4.p/P8.p do not divide,<br>some P5.p centering | EMS | 4:882874 | T/C | g02936 | *egl-20* / *Wnt* | Missense variant | Dichtel et al. 2001 |
| *iov-1* | *mf86* | Hypoinduction | TMP-UV | 39:154942-155133 | 191 bp deletion | g12432 | *lin-3* | Cis-regulatory deletion | Dichtel-Danjoy & Félix 2004 |
| *iov-2* | *mf76* | Hyperinduction 2° | TMP-UV | 10:74243-74877 | 634 bp deletion | g05993 | *smp-1* / *semaphorin* | Putative null | Dichtel-Danjoy & Félix 2004 |
| *iov-3* | *sy447* | Hyperinduction 2° | EMS | 86:24647 | A/T | g14741 | *plx-1* / *plexin* | Missense variant | Dichtel-Danjoy & Félix 2004 |
| *iov-3* | *mf52* | Hyperinduction 2° | EMS | 86:24066 | A/T | g14741 | *plx-1* / *plexin* | Missense variant | Dichtel-Danjoy & Félix 2004 |
| *iov-3* | *mf78* | Hyperinduction 2° | EMS | 86:27580-27583 | deletion | g14741 | *plx-1* / *plexin* | Premature stop | Dichtel-Danjoy & Félix 2004 |

\*The localization corresponds to the genomic position (**scaffold**: base pair). All molecular lesions in the table were identified by the mapping-by-sequencing approach, except the additional alleles *mf78* and *sy451* that were identified by PCR and Sanger sequencing of the gene.

**Table 2.** Quantification of large interspaces (gaps) between VPCs in *C. elegans* and *O. tipulae* plexin mutants, as determined by MH27 staining.

| Species | Strain | Phenotype | # animals |
| --- | --- | --- | --- |
| *C. elegans* | N2 | WT | > 50 |
| *C. elegans* | ST54: *plx-1(nc37)* | WT | 37 |
|  |  | Gap P7.p - P8.p | 13 |
|  |  | Gap P6.p - P7.p | 6 |
|  |  | Gaps P4.p - P5.p and P6.p - P7.p | 1 |
| *O. tipulae* | CEW1 | WT | > 30 |
| *O. tipulae* | JU108: *Oti-plx-1(mf78)* | WT | 35 |
|  |  | Gap P4.p - P5.p | 2 |
|  |  | Gaps P4.p - P5.p and P7.p - P8.p | 1 |

## Acknowledgements

We thank Aurélien Richaud for performing a smFISH experiment, Joao Picao Osorio for comments on the manuscript, Michalis Barkoulas for his advice on the *Oscheius* smFISH, Mark Blaxter’s lab for their continuous support of nematode genomics as well as for maintaining the *Caenorhabditis* website (http://www.caenorhabditis.org/), Benjamin Podbilewicz for advice on the immunofluorescence experiments, and Sana Dieudonné, Aurélien Richaud and Clément Dubois for advice and help in the use of the mapping by sequencing technique. Some strains were provided by the CGC, which is funded by NIH Office of Research Infrastructure Programs (P40 OD010440). We thank Wormbase.

## Author contributions

MAF, FB and AMVV designed the experiments. MAF and FB performed crosses, isolated DNA and analyzed the sequences, with input from AMVV. FB, MAF and AMVV identified *O. tipulae* gene homologs. AMVV performed the smFISH, CRISPR and DIC analyses. AMVV and MAF wrote the manuscript, with input from FB.

---

**Figure S1.**
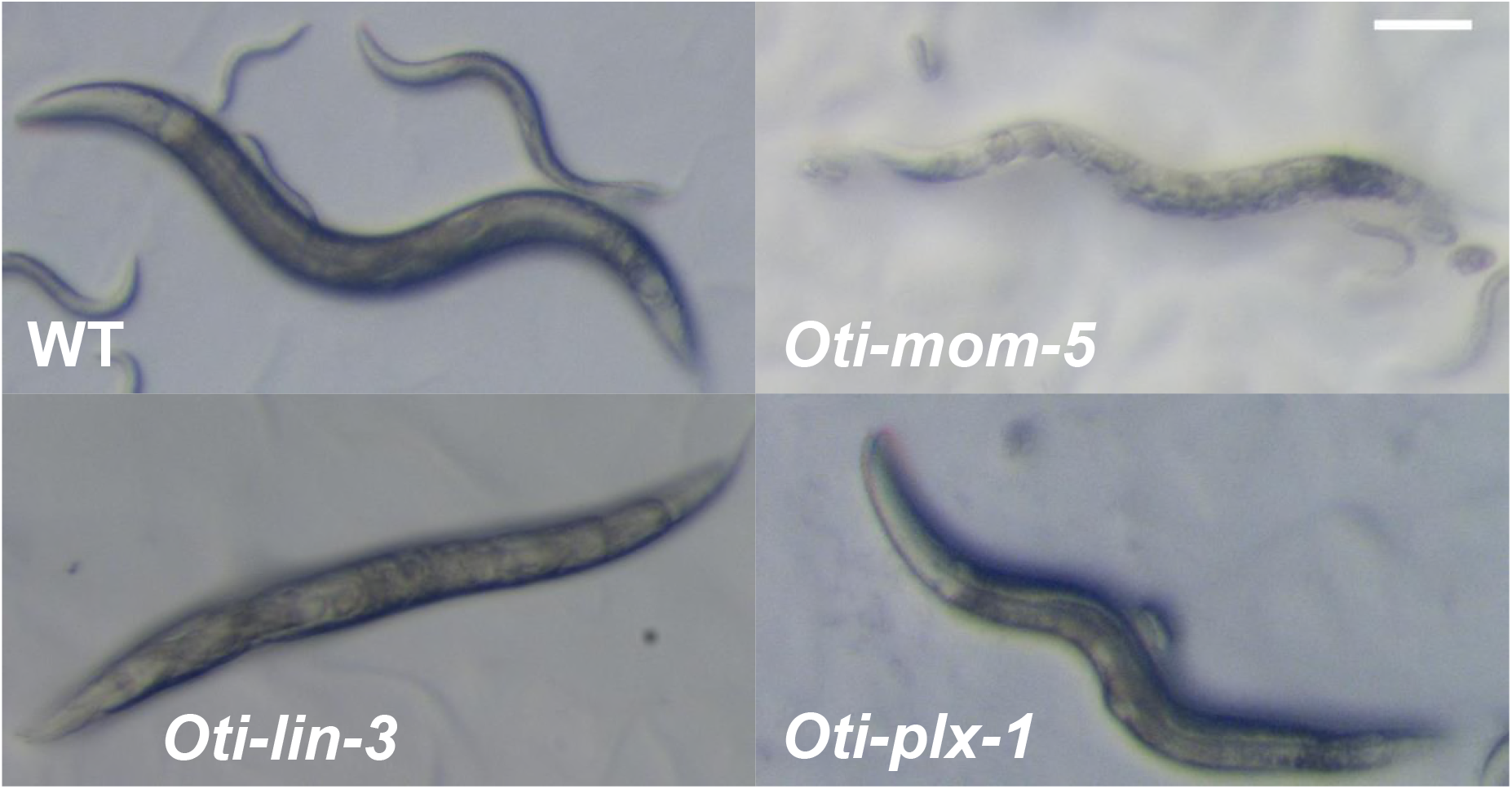
Dissecting microscope pictures of wild-type and mutant *O. tipulae* adult hermaphrodites. WT: wild-type. *Oti-lin-3(mf86)*: fully-penetrant egg-laying defective, forming a bag of worms. *Oti-mom-5(sy493)*: also partially egg-laying defective and protruding vulva. *Oti-plx-1(mf78)*: protruding vulva. All the images are set to the same scale. Scale bar: 100 micrometers.

**Figure S2.**
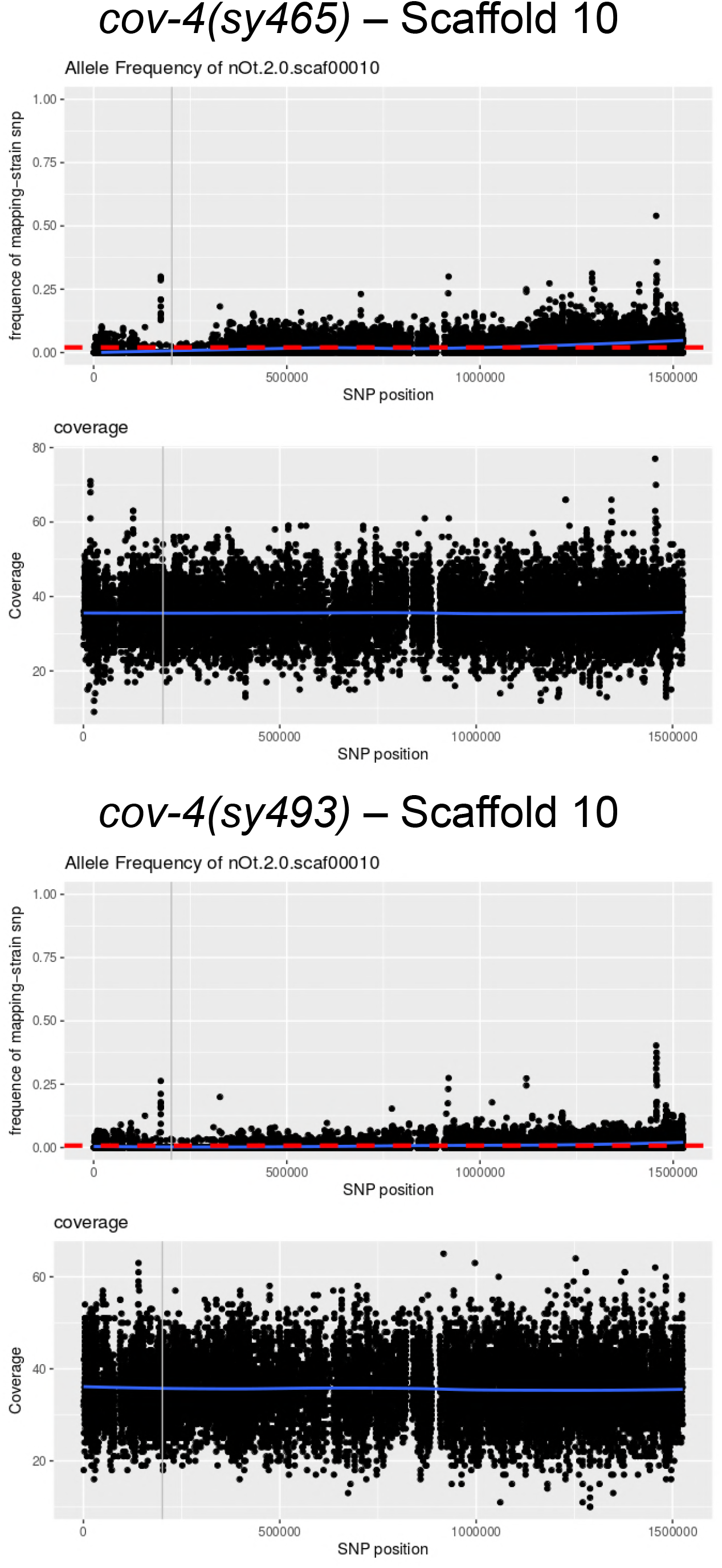
Example of mapping-by-sequencing in *O. tipulae*. Graphs showing the frequency of JU170 calls for single-nucleotide polymorphisms between CEW1 (reference wild isolate, on which the mutagenesis was conducted) and JU170 (alternative wild isolate used for mapping) along scaffold 10 of genome assembly nOt.2.0, for two alleles of the *cov-4* locus, called *sy465* and *sy493*. The location of *Oti-mom-5* is marked by a grey line. See (Besnard *et al*. 2017) for further details.

**Figure S3.**
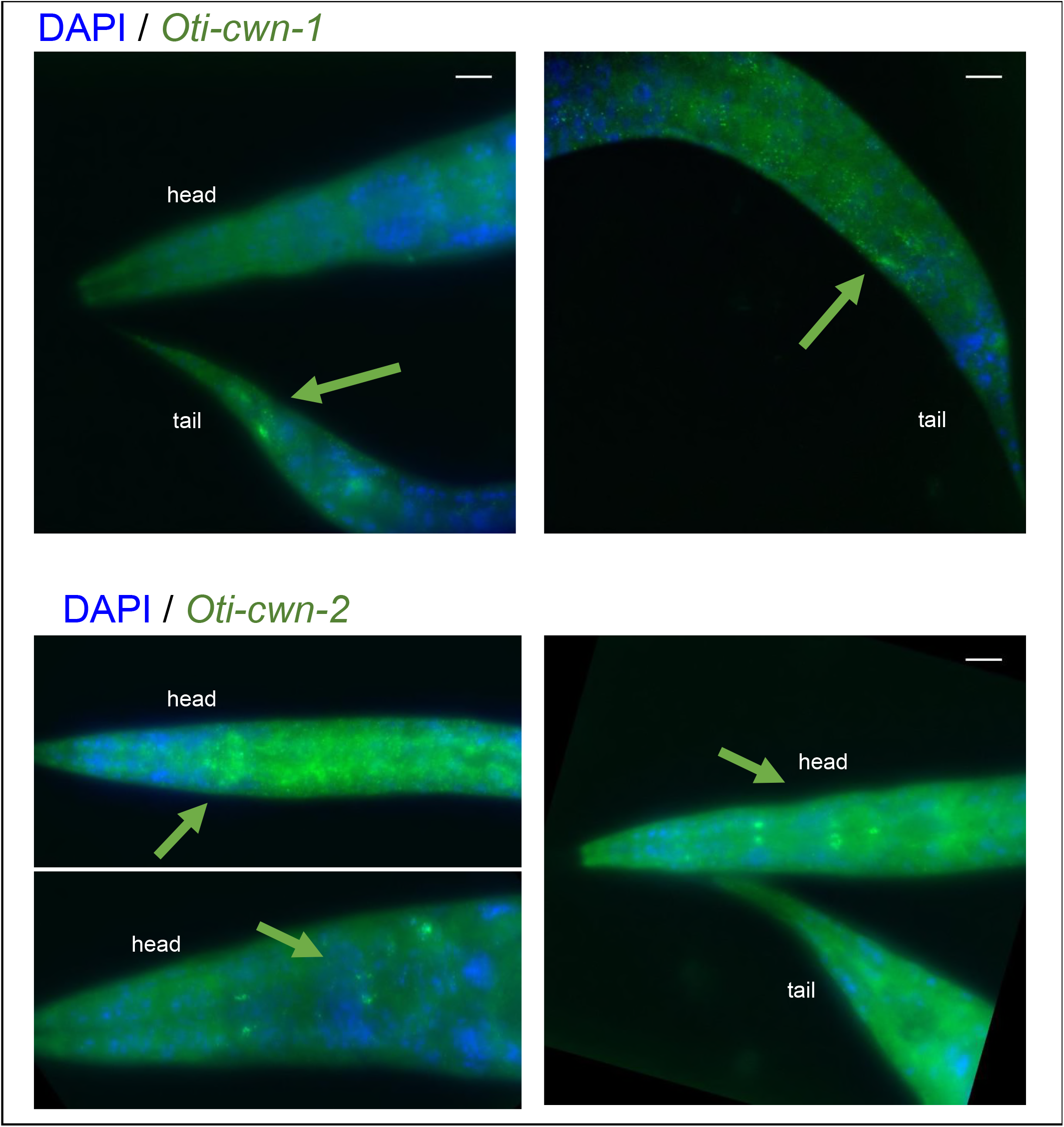
Single-molecule FISH of Wnt genes *Oti-cwn-1* and *Oti-cwn-2*. Wnt mRNAs are visible as green dots. The animals were also labeled with DAPI (in blue), labeling nuclei. *Oti-cwn-1* is visible only at the posterior part of the animal (green arrows), while *Oti-cwn-2* mRNAs (green arrows) appear in the pharynx and the anterior part of the animal. All the images are set to the same scale. The size of the bar is 10 micrometers.

**Figure S4.**
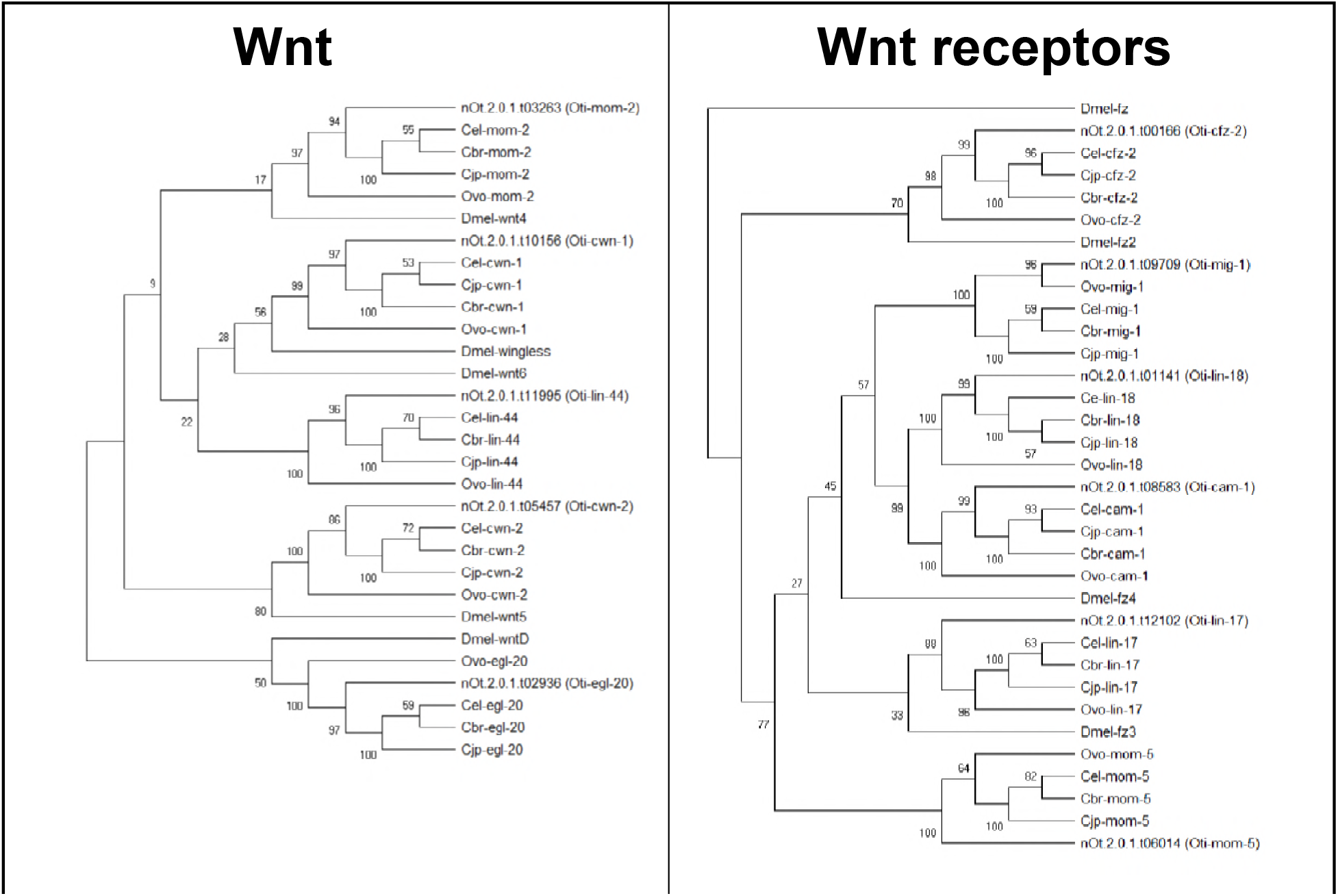
Phylogenetic relationship between Wnt genes inside and outside the *Caenorhabditis* clade. The cladograms were inferred using the Neighbor-Joining method with 1000 replicates for boostrapping. The percentage of replicate trees in which the associated taxa clustered together in the bootstrap test are shown next to the branches. Evolutionary analyses were conducted in MEGA X. Abbreviations: Cbr (C. *briggsae)*, Cel (C. *elegans*), Cjp (C. *japonica*), Dmel *(Drosophila melagonaster)*, Oti (O. *tipulae*), Ovo *(Onchocerca volvulus)*.

**Figure S5.**
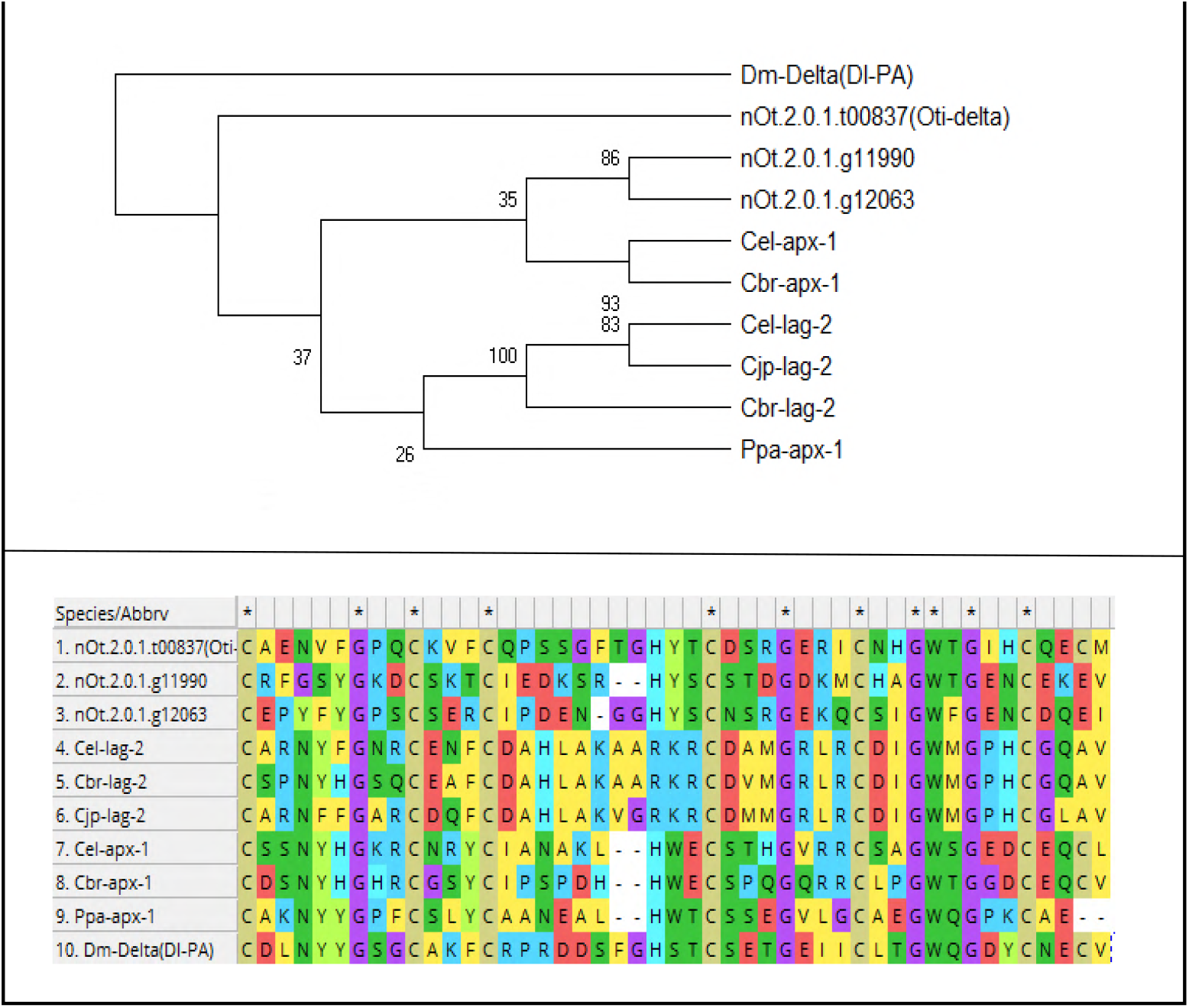
Identification and phylogentic relationships of Delta/Serrate/Lag-2 (DSL) proteins in *O. tipulae*. **Top panel:** Cladogram inferred using the Neighbor-Joining method with 1000 replicates for boostrapping. Abbreviations: Cbr (C. *briggsae*), Cel (C. *elegans*), Cjp (C. *japonica*), Dm *(Drosophila melagonaster)*, Oti (O. *tipulae*), Ppa *(Pristionchus pacificus)*. **Bottom panel:** Alignement of the delta motif used to calculate the molecular distances between DSL proteins.

**Figure S6.**
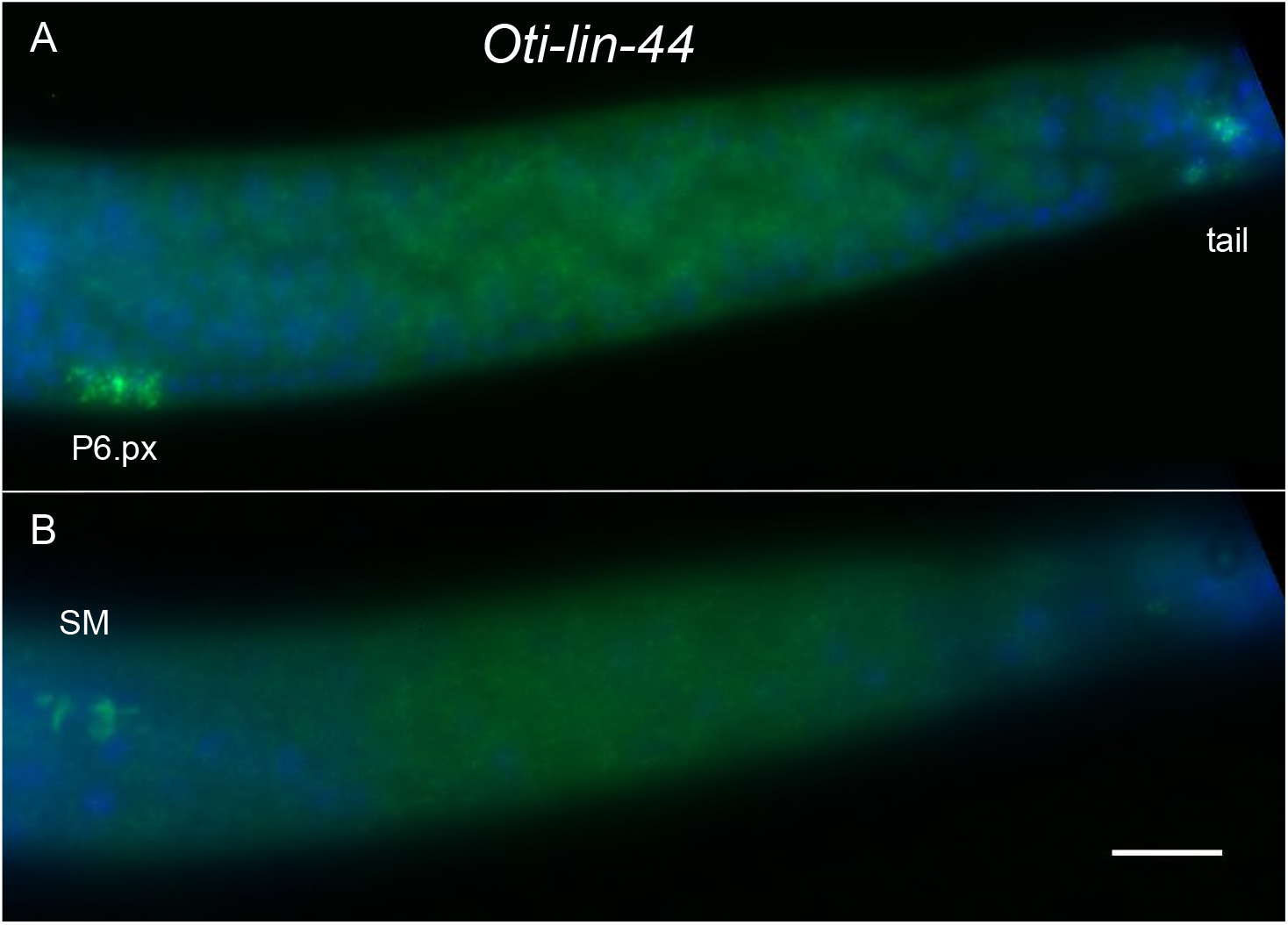
Expression profile of *Oti-lin-44* revealed by smFISH. L3 stage larva of *O. tipulae* CEW1 larva in two focal planes. (A) Staining is visible in the tail and the daughters of P6.p in the mid-focal plane. (B) Staining is visible in the cytoplasm of a sex myoblast in a lateral focal plane.

**Figure S7.**
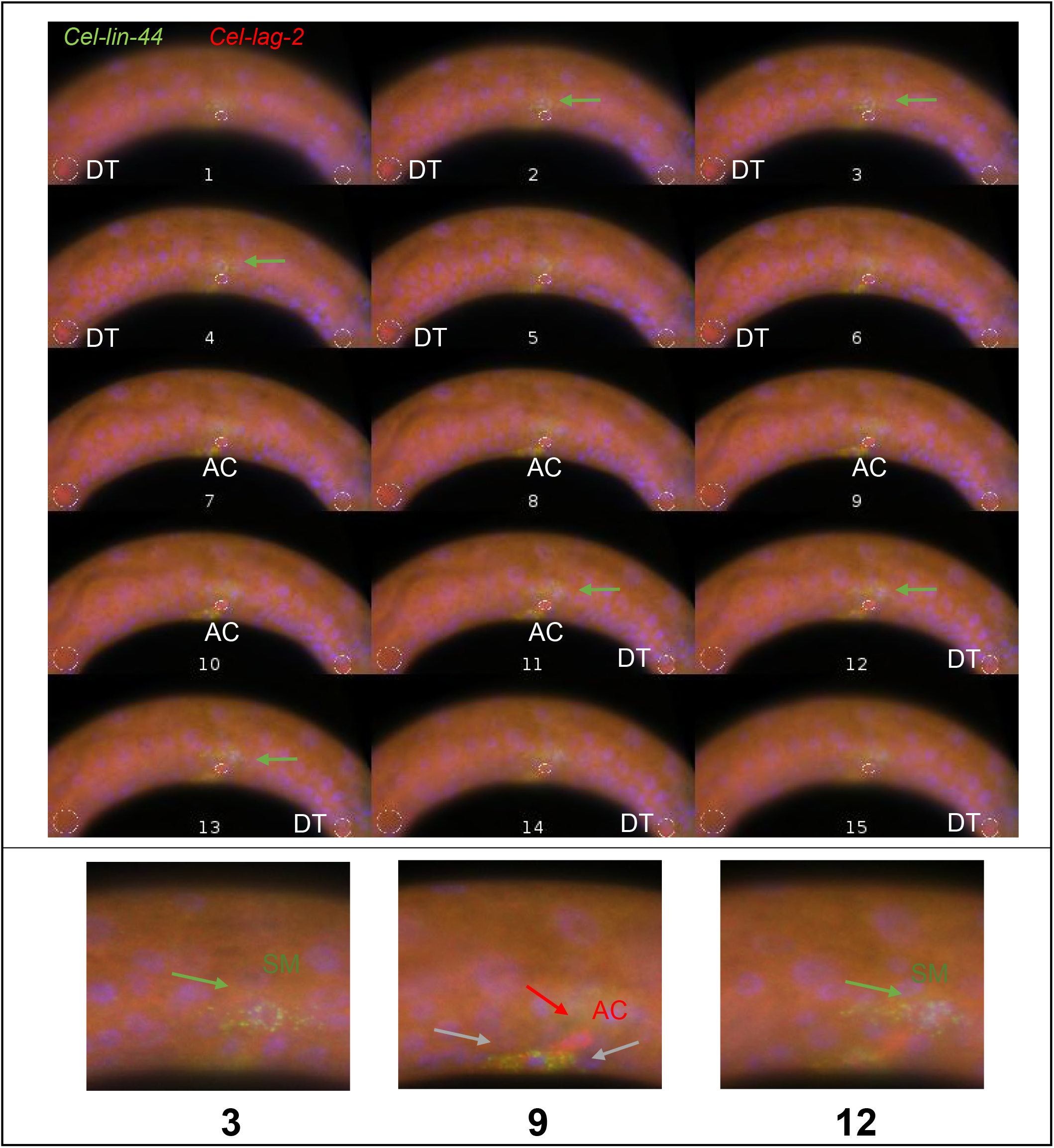
Expression profile of *Cel-lin-44* revealed by smFISH. **Top panel:** Z cuts of a L3 stage *C. elegans* N2 larva labelled with DAPI (blue) and smFISH probes for *lag-2* (red) and for *lin-44* (green). Each image is separated by 0.7 microns. The images were annotated when certain features were in focus, i.e. left/right distal tip cells (DT) - anchor cell (AC). **Bottom panel:** Enlarged set of images revealing the expression of *lin-44* in the sex myoblast (SM, green arrows) and the daugthers of P6.p (grey arrow), but not in the AC (red arrow).

**Table S1.** List of strains used in this study.

**Table S2.** Sequences of DNA primers used in this study. Sequencing primers to verify by Sanger sequencing the mutations identified by the mapping by sequencing approach, and to identify the molecular lesion in additional alleles.

**Table S3.** Sequences of smFISH probes used in this study. The fluorophore coupled to each probe is noted at the end of the set name.

**Table S4.** smFISH quantifications, distance measurements and vulval cell fates used in this study.

